# The canonical single-stranded DNA binding protein SSB is not an essential replication protein but an RNA chaperon in the hyperthermophilic archaeon *Saccharolobus islandicus* REY15A

**DOI:** 10.1101/2023.07.06.548054

**Authors:** Yuanxi Xiao, Zhichao Jiang, Mengqi Zhang, Xuemei Zhang, Qi Gan, Yunfeng Yang, Pengju Wu, Xu Feng, Jinfeng Ni, Xiuzhu Dong, Qunxin She, Qihong Huang, Yulong Shen

**Affiliations:** CRISPR and Archaea Biology Research Center, State Key Laboratory of Microbial Technology, Microbial Technology Institute, Shandong University, 266237, Qingdao, China; State Key Laboratory of Microbial Resources, Institute of Microbiology, Chinese Academy of Sciences, Beijing, China

**Keywords:** single-stranded DNA binding protein (SSB), archaea, Sulfolobales, cold shock protein, RNA chaperon

## Abstract

Single-stranded DNA binding proteins (SSBs) have been regarded as indispensable factors in all three domains of life since they play vital roles in DNA replication. Herein, we report that genes coding for the canonical SSB (SisSSB) and the non-canonical SSB (SisDBP) in the hyperthermophilic archaeon *Saccharolobus islandicus* REY15A can both be deleted. The growth is not affected, and the cell cycle progression and genome stability of the deletion strains is not impaired, suggesting that SisSSB and SisDBP are not essential for cell viability. Interestingly, at a lower temperature (55°C), the protein level of SisSSB increases ∼1.8 fold in the wild type and the growth of Δ*Sisssb* and Δ*Sisssb*Δ*Sisdbp* is retarded. SisSSB exhibits melting activity on dsRNA and DNA/RNA hybrid *in vitro* and unwinding RNA hairpin in *Escherichia coli*. Furthermore, the core SisSSB domain is able to complement the absence of the cold shock proteins CspABGE in *E. coli*, suggesting that SisSSB functions as RNA chaperon. We show that a two-fold overexpression of SisSSB is beneficial to the cell growth at lower temperature, but it has detrimental effect on the cell growth and cell cycle progression at normal growth temperature, which differs from bacterial Csp proteins. Importantly, these *in vitro* and *in vivo* activities are conserved in SSB subtype SSB-1 in Crenarchaeota species that lack bacterial Csp homologs. Overall, we have clarified the function of the archaeal canonical SSB which does not function as a DNA processing factor, but plays a role in processes requiring dsRNA or DNA/RNA hybrid unwinding.

## Introduction

Single-stranded DNA binding proteins (SSBs) are conserved and ubiquitous in bacteria, archaea, and eukaryotes. SSBs play vital roles in DNA replication, homologous recombination, and DNA damage repair (1–3). Canonical SSBs share a structural fold called oligosaccharide/oligonucleotide-binding (OB) fold that is responsible for ssDNA binding (4). In the replisome, SSBs protect ssDNA from degradation and prevent secondary structure formation by binding to ssDNA. SSBs also recruit replication proteins for the initiation and elongation of DNA replication(5–7). SSBs interact with a variety of proteins for DNA metabolism and are believed to play irreplaceable roles in genomic stability maintenance (8–10).

In eukaryotes, the major single-stranded DNA binding factor is the heterotrimeric replication protein A (RPA), which is made up of three subunits RPA1, RPA2, and RPA3, and contains six OB-folds in total (11). Four of the six OB-folds (DBD-A, DBD-B, DBD-C, and DBD-D) are responsible for ssDNA binding, while DBD-E is the structural center of RPA heterotrimeric and DBD-F works for protein-protein interaction (12). In bacteria, the *Escherichia coli* SSB (EcoSSB) comprises an OB-fold at the N-terminal for ssDNA binding and a disordered C-terminal tail that mediates protein-protein interactions, and it always forms a tetramer in solution (13). Whereas the archaeal single-stranded DNA binding proteins are diverse. Some archaeal species, those of Euryarchaeota in particular, such as *Haloferax volcanii* (14) and *Pyrococcus furiosus* (15), encode eukaryotic RPA-like heterotrimer single-strands DNA binding proteins. Many crenarchaeal SSB, for example, SSB from *Saccharolobus solfataricus* (formerly *Sulfolobus solfataricus*), SsoSSB, comprises an N-terminal OB-fold domain and a flexible C-terminal tail like EcoSSB. However, the N-terminal OB-fold structure resembles more to that of eukaryotic RPA DBD-B domain than the EcoSSB OB-fold domain (16). Further, it was found that Thermoproteales of the phylum Crenarchaeaota lack any recognizable canonical SSB but possess a non-canonical SSB called ThermoDBP. It was thus assumed that ThermDBP displaces the canonical SSB for ssDNA binding (17).

The model crenarchaeon *Saccharolobus islandicus* (a close relative of *S. sofataricus*) encodes both SsoSSB and ThermoDBP homologs, which we named as SisSSB (SiRe_0161) and SisDBP (SiRe_1003) (18,19). *Sisssb* was identified as an essential gene in *S. islandicu*s (20). Surprisingly, different from RPA in human, the gene coding for the canonical SSB could be deleted in *Sulfolobus acidocaldarius* (21). These raise interesting questions such as whether the crenarchaeal DBP is able to function as a replication factor and whether the canonical SSB could have a different function. In addition, the interactome of canonical SSB is poorly understood in archaea. In short, the exact physiological role of crenarchaeal SSBs remains ambiguous and unclear.

In this study, we performed genetic, biochemical and transcriptomic studies on both the canonical SSB and ThermoDBP in the hyperthermophilic archaeon *S. islandicus* REY15A (17,22). We found that either *ssb* or *dbp* or both can be deleted. The deletion did not cause apparent physiological changes nor the genome instability, but led to growth retardance at lower temperatures. A two-fold overexpression of SisSSB resulted in overall elongation of the cell cycle, and high overexpression was lethal to the cell. SisSSB was further determined to function as a cold shock protein in *S. islandicus*. As SisSSB exhibited RNA unwinding and anti-transcriptional termination abilities, we conclude that the canonical single-stranded DNA binding protein in *S. islandicus* REY15A is not an essential replication factor but functions as an RNA chaperon and cold shock protein.

## Results

### *Sisssb* and *Sisdbp* can be deleted in *S. islandicus* REY15A

*S. islandicus* REY15A contains two annotated single-stranded DNA binding proteins, the canonical SSB that contains an OB-fold (SisSSB) and the non-canonical Thermoproteales DBP (ThermoDBP) homolog (SisDBP)(16–19). SSB and RPA have long been known as essential replication factors in bacteria and eukaryotes, respectively (2,6,23,24). Although the OB-fold structure of the crenarchaeal SSB resembles more the eukaryotic RPA OB fold, the role of *Sulfolobus* SSB in DNA replication has never been verified. Recently, the gene encoding SSB was reported to be dispensable in *Sulfolobus acidocaldarius* (21), which implies that *ssb* is not an essential gene in archaea. To address whether the SisDBP was able to complement the loss of *ssb* and what is the real function of SSB, we performed genetic analysis on the two single stranded-DNA binding proteins. Using the endogenous CRISPR-Cas based method (25), we firstly obtained Δ*ssb* and Δ*dbp* (Figure S1A). To our surprise, the strain Δ*dbp*Δ*ssb* with double deletion of the two genes was also obtained by transforming the pGE-*dbp*-Knockout plasmid into Δ*ssb* cells and subsequent mutant screening. The deletion was confirmed by PCR and Western blotting (Figure 1A and 1B). Intriguingly, the growth of the deletion strains at 75℃ in the liquid STVU medium did not show apparent difference (Figure 1C), in fact, the cell density of strains lacking *ssb* even increased slightly (Figure 1C). We found that the cell morphology was the same as the control E233S (Figure S2F).

**Figure 1.**
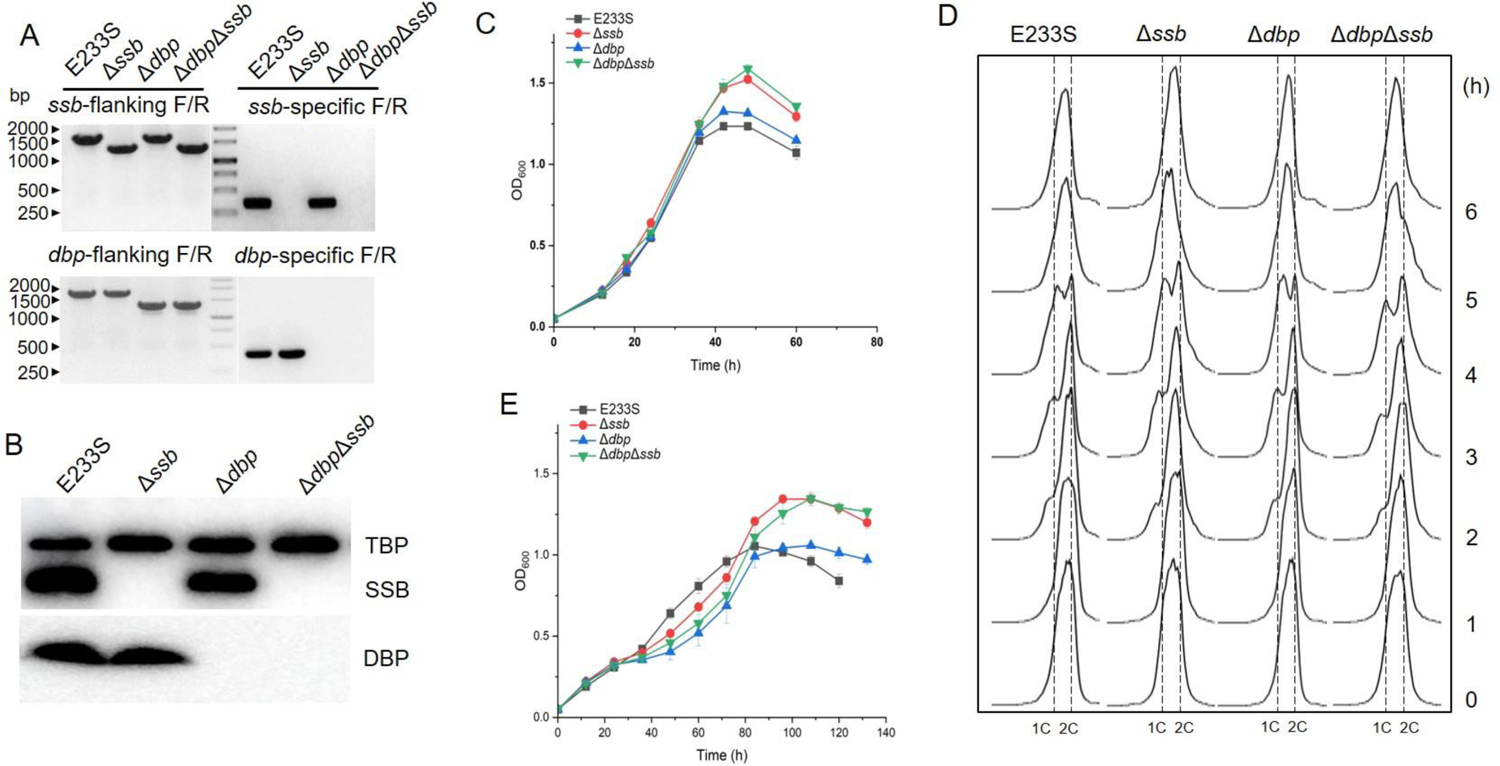
Growth and flow cytometry analysis of the knock-out strains of *Sisssb* and *Sisdbp*. (**A**) PCR verification of the knock-out strains Δ*ssb*, Δ*dbp* and Δ*dbp*Δ*ssb*. Genomic DNA and two primer pairs, flanking F/R and gene specific F/R were used for the analysis. (**B**) Western blotting analysis of the knock-out strains using whole cell lysate. Cells were collected at OD_600_=0.5∼0.8, disrupted by sonication. Primary antibody of SisDBP was added alone, while those of SisTBP and SisSSB were incubated simultaneously. Anti-SisTBP was used as the loading control. (**C**) Growth curves under normal conditions. Cells were cultured with shaking at 110 rpm and 75℃ in STVU medium with an initial OD_600_ 0.05. The values were calculated based on three biological repeats. (**D**) Flow cytometry analysis of the knock-out strains. Acetic acid (6 mM) was added into the culture when the OD_600_ reached 0.15∼0.2. Samples were taken at different time (0, 1, 2, 3, 4, 5, 6, 7, and 8 h) and analyzed as described in the Materials and Methods. 1C or 2C indicate one or two copies of chromosomes. (**E**) Growth curves of cells treated with 4-NQO. Cells were cultured as in (**C**) except that 4-NQO (3 μM) was added into the culture when the OD_600_ reached 0.2 (at 12h). The values were calculated based on three biological repeats.

### Deletion of the *ssb* does not impair the cell cycle progression and sensitivity to DNA damage agents

According to previous reports, ssDNA needs to be protected during DNA replication (5,6). We employed flow cytometry to check whether the S phase (DNA synthesis) was affected or not. Cells were synchronized at G2 phase (Figure 1D, 0 h), then the cell cycle was restarted by removing the acetic acid in the medium following a method as described in our previous report (26). The flow cytometry profiles showed that the wild type E233S exhibited an obvious peak of 1C (cells with 1 copy of chromosome) at 2 h, and the same 1C peaks also appeared at 2 h in Δ*ssb*, Δ*dbp*, and Δ*dbp*Δ*ssb* cells. It took about 3 h for the appearance of two copies of chromosomes (2C) peak (from 2h to 5 h) as the wild type. It seems that deletion of either *ssb*, *dbp*, or both did not affect the DNA replication efficiency.

It is generally accepted that ssDNA is generated after DNA damage and is involved in DNA damage response (DDR) and DNA repair (27–29). To check if SSB and DBP are involved in DNA repair, the growth of the deletion strains was followed by measuring OD of their cultures in the presence of each of the four different DNA damage agents, including 4-nitroquinoline N-oxide (4-NQO), cisplatin, hydroxyurea (HU), and methyl-methanesulfonate (MMS), which induce DNA bulky adduct, intra-strand crosslink, stalling of the DNA replication fork, and nucleotide alkylating, respectively. We found that these DNA damage agents showed no apparent detrimental effect on the growth of these deletion strains compared with E233S (Figure 1E and Figure S2A-2C).

In *S. islandicus* REY15A, a DDR network centered by Orc1-2 plays an important role in response to and repair of damaged DNA (30,31). In the presence of DNA damage agent NQO or UV treatment, Orc1-2 up-regulates the expression of genes coding for components of pili, DNA exchange system (Ced), and the homologous recombination repair (HRR) systems. When *S. islandicus* cells encounters extensive DNA damage, they form aggregates in which intercellular DNA exchange is believed to occur, providing undamaged DNA templates for HRR in recipient cells. To explore whether SSB and DBP in *S. islandicus* are involved in DNA damage signaling, cell morphology and cell aggregation were examined. As shown in Figure S2D-E, cell aggregation occurred more rapidly in Δ*ssb* than in E233S. Three hours after NQO treatment, 65% of Δ*ssb* cells formed aggregation, while almost no cell aggregation was observed in E233S. In addition, 12h after NQO treatment, Δ*ssb* exhibited higher cell aggregation than E233S (89% vs 59%). During DDR, Δ*ssb* displayed a higher cell aggregation ratio than E233S. Nevertheless, cell aggregation in Δ*ssb*, same as E233S, decreased gradually after 12h, and finally disappeared. The results suggested that the DNA damage response in Δ*ssb* is more efficient than that of the wild type. The deletion of DBP did not increase the cell aggregation ratio after NQO treatment. These results indicate that the two ssDNA binding proteins are not directly involved in DNA damage response and repair, but play other roles independent of DNA replication, DNA damage response, and DNA repair. The exact mechanism underlying *ssb* deletion-induced quicker and more efficient DDR response remains unknown, however, loss of SSB may make more *ssDNA* exposed which could quicker initiate DDR.

### Elevated SSB is found at a lower growth temperature

In *E. coli*, *ssb* mutant strain *ssb-1* is temperature-sensitive and displays lethal phenotype at 30℃ (32). In the thermophilic crenarchaeon *S. acidocaldarius, ssb* deletion caused retarded growth at lower temperatures (21). We then tested the growth of the *ssb* deletion and overexpression strains of *S. islandicus* REY15A at lower temperatures 65℃, 60℃ and 55℃. In agreement with the result in *S. acidocaldarius* (21), deletion of *ssb* or both of *dbp* and *ssb* led to much slower growth than strain E233S at 55℃, and the *ssb* knock-down strain *Para::ssb* in which the native *ssb* promoter is replaced with an arabinose-inducible promoter (*ParaS*-SD, see below for details), has the same growth with *ssb* deletion strain (Figure 2A and Figure S3A and S3B). As temperature decreased, the growth inhibition became more pronounced. However, in the presence 0.2% arabinose, the 2-fold SSB overexpression strain *Para::ssb* exhibited better growth than Δ*ssb* and the knock-down strains at 55℃, although it still grew slower than E233S (Figure S3). Our results are consistent with that in *S. acidocaldarious* (21). The results imply that the archaeal SSBs function in cold adaption.

**Figure 2.**
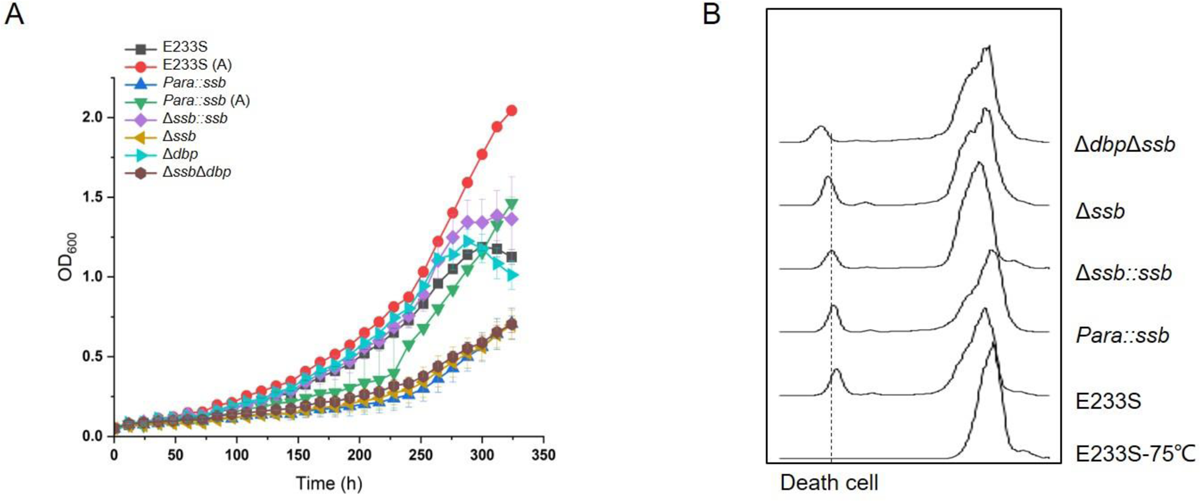
SSB-deficient cells exhibited slow growth at lower temperature. **(A)** Growth curves of SSB-deficient strains at 55℃. Cells were cultured in liquid STVU at 75℃ to OD_600_= 0.6∼0.8 and used as inoculates to be cultivated with an initial OD_600_ of 0.05 at 55℃ with shaking. The optical density was monitored and the values were calculated based on measurements of three biological replicates. **(B)** Cytometry profiles of the strains grown at 55℃. Samples were taken at 72 h and analyzed. Cells of E233S cultivated to middle logarithmic phase (OD_600_∼0.4) at 75℃ was used as a control.

The *E. coli* SSB has a much weaker affinity to ssRNA than ssDNA (33). Human RPA also has a high affinity for ssDNA and low affinity for ssRNA, the difference is at least three orders of magnitude (34). Previous studies showed that SSB from the hyperthermophilic crenarchaeon *S. solfataricus* binds to both ssDNA and ssRNA with almost equal affinities (22). This extraordinary nucleic acid binding feature of archaeal SSB also implies that they could have other unknown functions.

It is known that the levels of the bacterial cold shock proteins (Csps) increase in response to temperature downshift (35). Csps are supposed to function as RNA chaperones which prevent secondary structure formation of mRNAs so to facilitate translation at low temperature (36,37). Csps are required for bacteria to adapt to ambient temperatures, and deletion of Csps results in a significant growth deficiency at low temperature in bacteria (38). Given that deletion of *ssb* also caused a cold sensitivity phenotype of *S. islandicus*, we assume that SisSSB may function as RNA chaperon as the bacterial Csps. To test this hypothesis, we detected the protein level of SisSSB in E233S at 75℃ and 55℃, respectively. The cells were cultured in liquid STVU medium at 75℃ to middle logarithmic phase (OD_600_≈0.4), and then cultured at 55℃ for 24 h. As expected, the SisSSB protein level increased to 1.8 folds when the culture was shifted from 75℃ to 55℃ (Figure 3A and 3B). In addition, this higher level of SisSSB maintained for further 3 days at 55℃ (Figure 3C). These observations suggest that SSB play a role in cold shock response.

**Figure 3.**
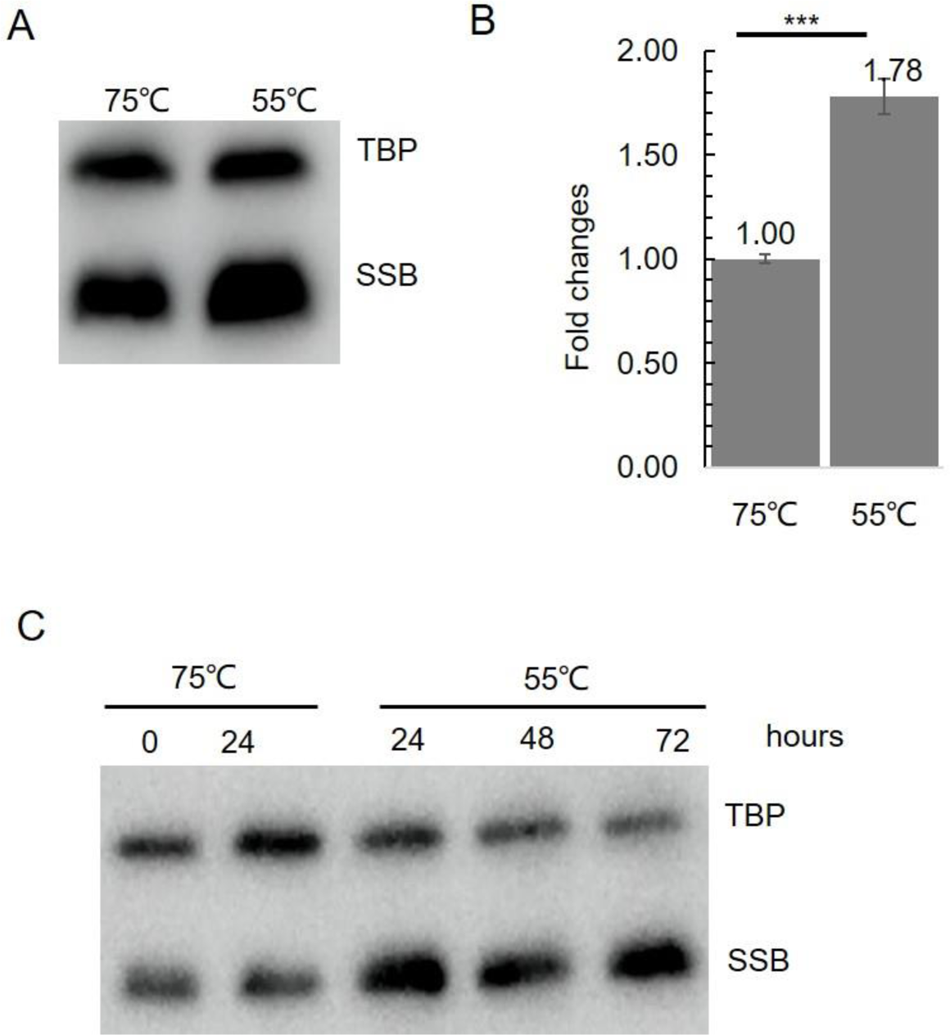
The expression level of SisSSB increases at lower temperature. **(A)** Western blotting analysis of SisSSB at 75℃ and 55℃. E233S cells were cultured at 75℃ to middle logarithmic phase (OD_600_∼0.4) then the cultures were moved to a shaker at 55℃. After cultivated for 24 h at 55℃, the cells were taken and disrupted by sonication. The cell lysates were subjected to SDS-PAGE and Western blotting analysis with the antibodies against SisSSB and SisTBP (loading control). **(B)** Quantitative analysis of the result in Fig (A). **(C)** Western blotting of SisSSB at 55℃ at different times. E233S cells were cultured at 75℃ to middle logarithmic phase (OD_600_∼0.4). The samples were taken and divided evenly into the culture flasks. One was incubated and cultured at 75℃ and the other was at 55℃ shaker. Cells were taken from the cultures after 24, 48, and 72 h, respectively.

### SisSSB has anti-transcription termination activities *in vivo* and can unwind dsRNA *in vitro*

To investigate if SSB has RNA chaperon activity *in vivo*, we used *E. coli* RL211 for the analysis (39,40). This strain harbors a chloramphenicol resistance gene (chloramphenicol acetyl transferase, CAT) as a reporter located downstream of a strong *trpL* terminator (41). A pINIII vector carrying IPTG-inducible SisSSB was transformed into RL211, and upon the terminator is melted, the CAT gene will be expressed, enabling the strain to grow in the presence of chloramphenicol. *E. coli* CspA, CspE and a confirmed RNA chaperone Sis10b in *S. islandicus* REY15A were used as positive controls. As shown in Figure 4A, same as CspA and CspE, cells carrying pIN-SisSSB grew well on chloramphenicol-containing agar plate even without IPTG induction, and SisSSB exhibited a stronger ability than Sis10b at 37℃.

**Figure 4.**
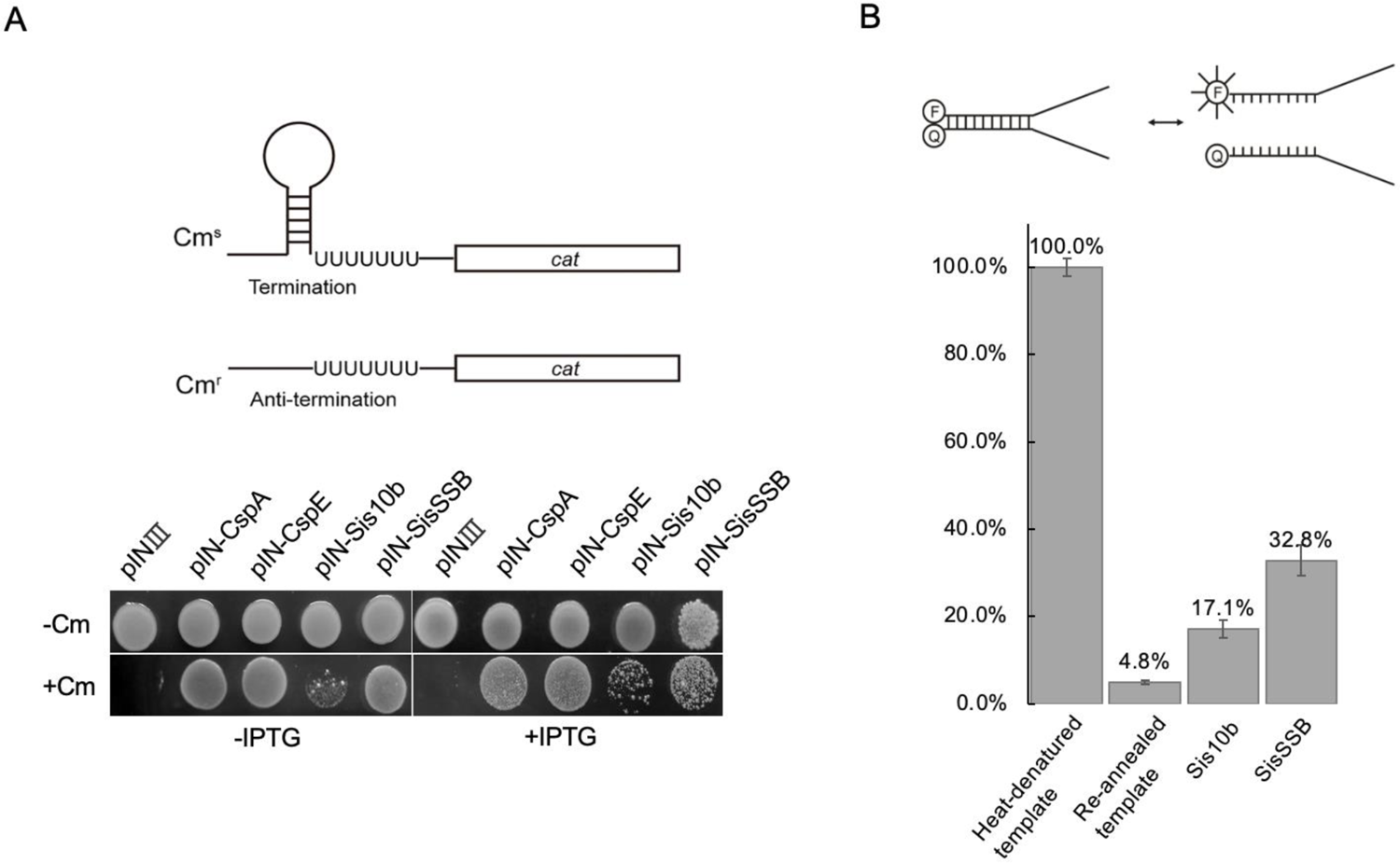
SisSSB has RNA chaperon activities *in vivo* and *in vitro.* (**A**) SisSSB has antitermination activity in *E. coli*. Upper panel, schematic of the *in vivo* antitermination assay using *E. coli* strain RL211. A chloramphenicol resistance gene *cat* cassette located downstream of the *trpL* terminators is used as a reporter. The hairpin RNA structure terminates the expression while its unwinding allows expression of the reporter. Lower panel, spot assay showing the antitermination activity of SisSSB. Cell were transformed with empty pINⅢ vector or pINⅢ carrying *cspA*, *cspE*, *Sis10b* or *Sisssb* were cultured and adjusted to an OD_600_ of 1.0 with the medium. Aliquot (6 μL) of each culture was plated onto a LB plate containing 100 μg/mL ampicillin, with or without 30 μg/mL chloramphenicol (+/−Cm) or 0.2 mM IPTG(+/-IPTG). **(B)** SisSSB is able to unwind dsRNA. Upper panel, schematic map showing the substrate and melted products. Lower panel, quantification of the dsRNA unwinding activity of SisSSB. The FAM-labeled ssRNA was defined as 100% and annealed dsRNA template was used as a negative control. Sis10b was used as a control.

These results suggest that SisSSB exhibited stronger anti-transcriptional termination ability than Sis10b *in vivo*. While with the addition of 0.2 mM IPTG, overexpression of SisSSB affected the growth of *E. coli* RL211 cells (Figure S4A and S4B).

To test if SisSSB has RNA chaperone activity *in vitro*, we used a molecular beacon assay (40). As shown in Figure 4B, a 40 nt 5’-FAM labelled ssRNA was annealed with another ssRNA labeled with a fluorescence quencher (BHQ1) at the 3’-end. The RNA chaperon Sis10b was used as a positive control. After annealing, the fluorescence of the partial duplex RNA decreased to ∼5% of that of the heat-denatured dsRNA substrate (Figure 4B). In the presence of SisSSB, the fluorescence intensity increased to ∼33% at 37°C which was higher than that of Sis10b (∼17%), consistent with the result of the anti-transcription termination assay *in vivo*. The above results demonstrate that SisSSB is able to unfold RNA *in vivo* in *E. coli* and *in vitro*.

### The OB fold domain of SisSSB can complement *E. coli* Csps

To further analyze the RNA chaperone activity of SisSSB, we used a cold sensitive *E. coli* strain BX04 in which the genes coding for the Csps (CspA, CspB, CspE and CspG) were all deleted (38,39). Plasmids pIN carrying IPTG-inducible CspE, Sis10b, SisSSB, or SisSSB mutants (Figure 5A) was transformed into BX04 and the transformants were grown in LB containing 100 μg/mL ampicillin. The cultures were diluted in gradience and spotted onto plates containing 100 μg/mL ampicillin with or without 0.2 mM IPTG and incubated for 3-5 days at 37℃ and 22℃, respectively. As shown in Figure 5B and Figure S5A, strain BX04 harboring CspE overexpression plasmid exhibited an optimal growth at 22℃, while BX04 with empty pIN plasmid was sensitive to low temperature. Interestingly, cells expressing SisSSB ΔC29 containing the OB-fold and its C-terminal structured segment showed similar growth to those expressing CspE at 22℃, while cells expressing SisSSB ΔC53 grew worse than those harboring CspE or SisSSBΔC29 plasmid but still better than those carrying the empty plasmid. Intriguingly, the growth of BX04 cells carrying plasmid overexpressing SisSSB(W56A), SisSSB(F79A), or SisSSB(W56AF79A) were impaired at 37℃ and 22℃, similar to that overexpressing the wild type SSB (Figure 5B and S5). EMSA assays showed that two C-truncated mutants ΔC53 and ΔC29 of SisSSB have nearly the same nucleic acid binding capacity as SisSSB WT (Figure S5B). Additionally, we analyzed the nucleotide binding activity of aromatic amino acid residue mutants W56A, F79A, and W56AF79A and found that the binding activities on ssDNA and ssRNA were all reduced (Figure S5B), in agreement with a previous report on SsoSSB (16). The results demonstrate that the N-terminal OB fold domain can functions as Csp to destabilize the RNA secondary structures at low temperature. We assume that the C-terminal flexible tail rich in negatively charged residues of SisSSB is harmful to *E. coli* and this effect seems to be independent of ssDNA binding,

**Figure 5.**
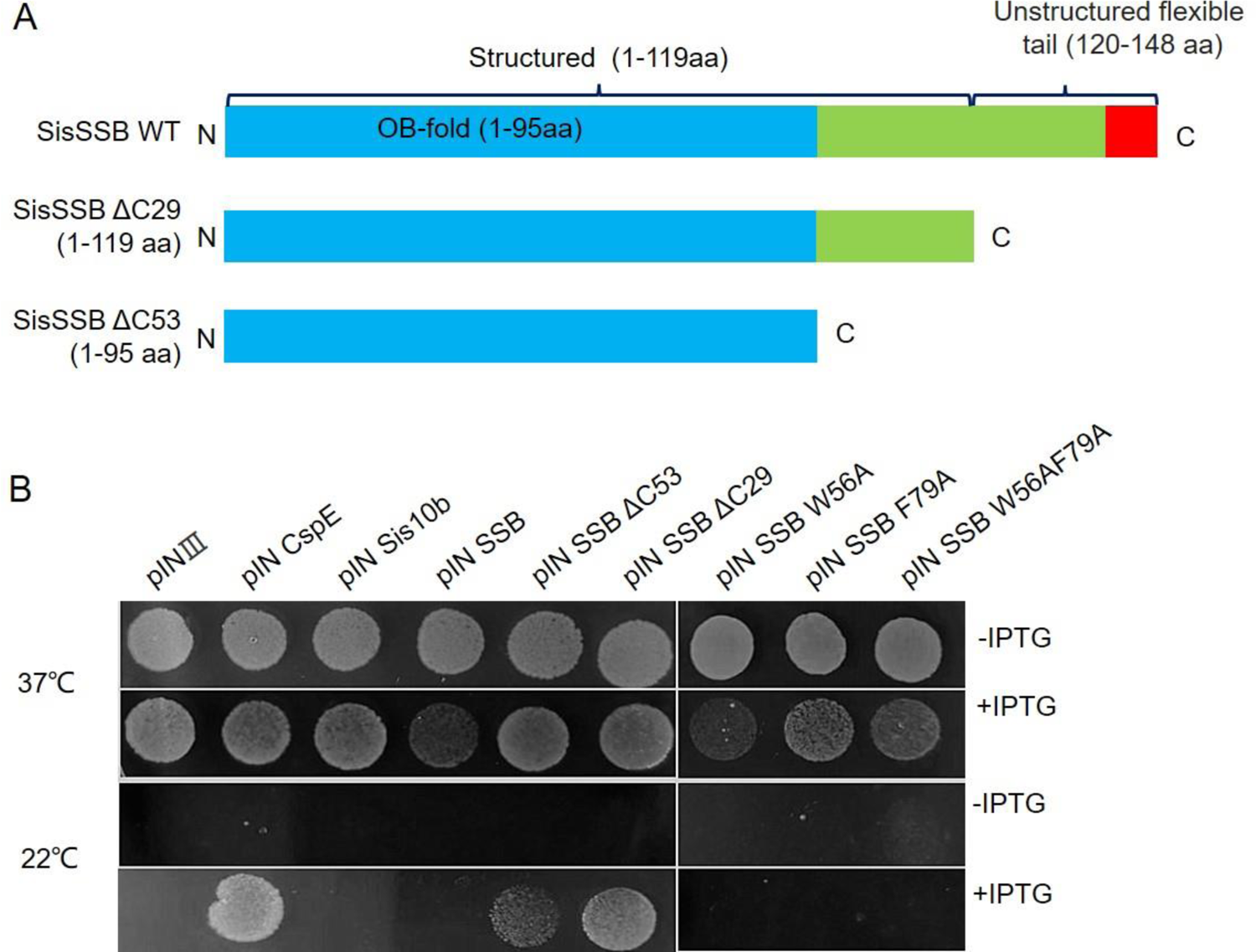
The OB-fold domain of SisSSB is able to complement the loss of *csp* gene in *E. coli.* (A) Schematic showing the domain structure of SisSSB and its C-terminal truncated mutants ΔC29 and ΔC53. The OB-fold was marked in blue and the region contain 8 acidic amino acids was marked in red. (B) *E. coli* BX04 carrying the pINIII plasmids for the expression of SisSSB and its C-terminal truncated or the binding site deficient mutants. Cells were cultured and adjusted to an OD_600_ of 1.0 with LB medium containing 100 μg/mL ampicillin and then serially diluted in 10 fold and spotted onto LB plates supplemented with ampicillin and with or without IPTG. The plates were incubated for 2–5 days at 37°C and 22°C, respectively.

### Overexpression of SisSSB leads to cell cycle elongation

To further understand the *in vivo* function of SisSSB, we attempted to overexpress SisSSB in *S. islandicus* REY15A. First, we constructed an overexpression plasmid pSeSD-SisSSB (no tag) and tried to get a transformant. We failed to obtain any colony after several attempts. Then, we successfully constructed a strain with the native promoter region of *Sisssb* (200 bp upstream of the start codon of *sire*_*0161*, Figure S1B) in the genome being replaced with an arabinose-inducible promoter (*ParaS*-SD) by the CRISPR-Cas-based genome editing method (25). The substitution was confirmed by PCR (Figure S1C) and Western blotting. As shown in Figure 6A and 6B, compared with E233S, the promoter substitution strain *Para::ssb* exhibited a low SSB expression (0.32 fold) in the sucrose-containing medium STVU and an elevated SSB protein level (2.11 fold) in the arabinose-containing medium compared to that in E233S. Interestingly, in the sucrose-containing medium, the strain grew almost the same as E233S (Figure S1D) at 75℃. However, in the arabinose-containing medium, the growth was significantly retardant (Figure 6C). The results indicate that although reduction of *ssb* expression has no apparent effect on the cell growth, the elevation of the expression as high as only about two folds has a drastic effect on the cell growth. Whereas overexpression of SSB using the pSeSD-based plasmid is lethal for the cells, perhaps due to even higher SSB expression. The results indicate that the function of SisSSB is different from that of EcoSSB in their corresponding cells.

**Figure 6.**
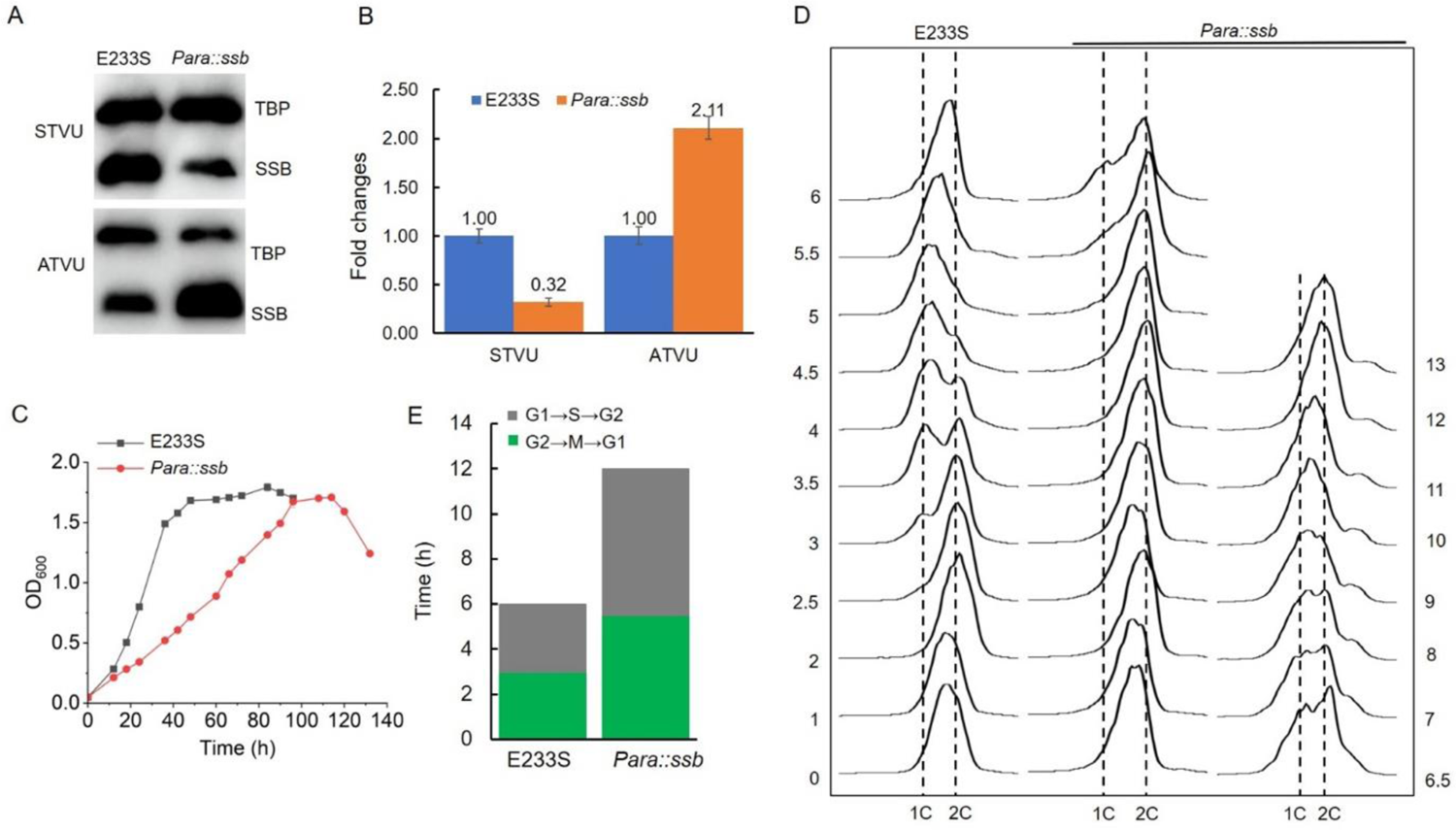
Overexpression of SisSSB has a global effect on the cell cycle progression and growth. (A) Assay of the expression levels in the promoter replacement strain under non-inducible and inducible conditions by Western blotting. Cells were cultured in liquid STVU or ATVU medium and collected at OD_600_=0.5∼0.8. The cells were then disrupted by sonication and the cell lysates were subjected to SDS-PAGE and Western blotting analysis with the antibodies against SisSSB and SisTBP. The antibodies against SisSSB and SisTBP were incubated at the same time. (B) Quantitative analysis of the result in (A). (C) Growth curves of the promoter substitution strain in comparison with the wild type E233S. The cells were cultured in liquid ATVU media with an initial OD_600_ 0.05 and with shaking (110 rpm) at 75℃. OD_600_ was monitored at 6 or 12 h interval. The values were based on measurements of three biological replicates. (D) Comparison of the cytometry profiles of the synchronized E233S and the promoter substitution. The cells were cultured in TSVU medium until the OD_600_ reached 0.15∼0.2 when acetic acid (6 mM) was added into the culture. After cultured for 3 h, 0.2% D-arabinose was added to induce the expression of SisSSB. After incubation for further 6 h, samples were taken at different time points (0, 1, 2, 2.5, 3, 3.5, 4, 4.5, 5, 5.5, 6, 6.5, 7, 8, 9, 10, 11, and 12 h) and subjected to the flow cytometry analysis. “1C” and “2C” indicate cells containing one and two copies of chromosomes, respectively. (E) Comparison of the cell cycle periods of E233S and SisSSB overexpression strain according to (D). G2→M→G1 (gray), from the cell cycle release to 1C appearance; G1→S→G2: 1C appearance to 2C peak.

In *E. coli*, it was reported that the SSB (EcoSSB) binds to ssDNA at the replication fork in different modes depending on the concentration of cations and the ratio of SSB/ssDNA (42,43). EcoSSB interacts with replication proteins during DNA replication, such as DNA polymerase χ subunit and primase in the replisome (44,45). We assumed that the growth inhibition caused by SisSSB overexpression might be due to the change of SSB/ssDNA ratio or the interaction with DNA replication proteins in the replisome, so that it may take a longer time for the DNA synthesis in the S period. To clarify this, cell cycle synchronization was performed to examine whether DNA replication was impaired under the elevated SSB level. As shown in Figure 6D, for the wild type, the peak of 1C was observed at 2.5 h, and it took about 3.5 h for the appearance of two copies of chromosomes (2C) peak (from 2.5 h to 6 h) in the arabinose medium. In contrast, the 1C peak in the SSB overexpression strain appeared at 5.5 h, which was about 2 h later than that of E233S. It took about 6.5 h from the appearance of 1C to that of 2C (from 5.5 to 12 h). These results show that although elongated S period (about two-time long) occurred in the overexpression strain, the whole cell cycle period also doubled (6 h:12 h) (Figure 6E). This suggests that SSB overexpression affects the cell cycle globally, not just the DNA synthesis. The above results reinforce that SSB is not an DNA replication factor and suggest that the RNA chaperon function of SSB is different from bacterial Csps, the expression of which is induced in great magnitude and overexpression of Csps does not has detrimental even lethal effect.

### Transcriptomic analysis of the *ssb* or *dbp* deletion strains and identification of the ssDNA interactome

To further probe the *in vivo* functions and relationship of SisSSB and SisDBP at transcription level, we performed transcriptomic analysis and analyzed the global gene expression changes caused by the deletion of *ssb* or *dbp*. Compared with E233S, 44 genes were up-regulated (>2 folds) in Δ*ssb*, while only 3 of them are related to DNA replication process after DNA damage (Table 1, Table S3). Three genes encoding SiRe_0614, SiRe_0615, and SiRe_0616 were up-regulated 2.27-3.25 folds after *ssb* deletion (Table 1). Among them, Dpo2 (SiRe_0615) has been identified as a functional eukaryotic pol zeta homolog responsible for DNA damage tolerance and plays an important role in archaeal DNA damage repair (46,47). SiRe_0614 and SiRe_0616 are well conserved in Sulfolobales genomes at the *dpo2* gene locus (48), they were supposed to code for Dpo2-associated factors (Daf) and work in concert with Dpo2 in DNA translesion synthesis (TLS) (46). Besides, 13 genes associated with DDR were also up-regulated, including Tfb3 (SiRe_1717) which is a transcription regulator of DDR, 3 genes involved in DNA transfer (SiRe_1316, SiRe_1879, SiRe_1881), and the rest 9 genes which are Orc1-2 dependent NQO-inducible genes (30). The rest 28 up-regulated genes and 18 down-regulated genes were mostly related to cell metabolism (Table S3). In contrast, only 3 genes were up-regulated and 5 genes were down-regulated in Δ*dbp* (Table S3), none of them seems to be involved in DNA metabolism. These data suggest that SisSSB is involved in the protection of ssDNA. Deletion of *ssb* may lead to more ssDNA generation which could ignite DNA damage response. On the other hand, the data on SisDBP do not give a clue to the function of SisDBP. In addition, it seems that deletion of *Sisssb* and *Sisdbp* did not affect each other at the transcription level, and transcription of either one is not affected by deletion of the other gene.

**Table 1.** Changes of the transcription levels of genes involved in DNA replication and DNA damage response due to *ssb* deletion

| Gene_ID | Fold change *<br>($\Delta ssb$ vs E233S) | Description |
| --- | --- | --- |
| DNA replication |  |  |
| SiRe_0614 | 3.26 | Uncharacterized protein associated with inactivated Dpo2 |
| SiRe_0615 | 2.31 | Dpo2 |
| SiRe_0616 | 2.27 | RecA/RadA recombinase, inactivated |
| Transcription factor |  |  |
| SiRe_1717 | 2.52 | TFB3 |
| DNA transfer |  |  |
| SiRe_1316 | 2.20 | CedA1 |
| SiRe_1879 | 2.48 | UpsE |
| SiRe_1881 | 4.79 | UpsA |
| Putative DDR related |  |  |
| SiRe_0020 | 2.34 | Uncharacterized protein |
| SiRe_0137 | 4.94 | MFS family permease |
| SiRe_0187 | 4.08 | Uncharacterized protein |
| SiRe_0589 | 3.47 | Adenine-specific DNA methylase containing a Zn-ribbon |
| SiRe_0670 | 8.67 | SWIM Zn-finger |
| SiRe_0269 | 3.85 | Uncharacterized protein |
| SiRe_1957 | 3.76 | Uncharacterized protein |
| SiRe_2100 | 6.09 | Membrane protein involved in DNA uptake |
| SiRe_2101 | 2.01 | Acyl-CoA dehydrogenase |

\*Genes with $\geq 2$ fold change and p-value $\leq 0.05$ were listed. The ratios were based on the data from three independent repeats.

Since none of the proteins was found capable of complement the DNA binding function of SSB through transcriptomic analysis, we then attempted to identify putative ssDNA binding proteins by *in vitro* pulldown using ssDNA as a bait and subsequent mass spectrometric (MS) analysis. We followed a strategy reported by Paytubi *et al* (17). A 5’-biotinylated 45 nt ssDNA was used as the bait. The ssDNA was bound to the agarose beads via the biotin-streptavidin interaction. The beads were incubated with the cell extracts of E233S, Δ*ssb* and Δ*dbp*Δ*ssb* for 2 h at 55°C to prey the putative ssDNA interacting proteins. The pull-down proteins were identified by Western blotting and MS analysis. As shown in Figure 7A, SisSSB was a dominant protein in the ssDNA interactome from the E233S cell lysate. Unexpectedly, in the samples of Δ*ssb* and Δ*dbp*Δ*ssb*, there is no protein that had similar abundance as SSB pulled with ssDNA, although SisDBP appeared to increase in the sample of Δ*ssb*. In addition to SisSSB and SisDBP, NusA (SiRe_1772) and an ATP-cone containing protein (SiRe_2062) were also identified in all the samples, but not at a comparable level with that of SisSSB (Figure 7A). As shown in Table S4, SisSSB had the highest score from sample of E233S in the identified proteins, while SiRe_1772 and SiRe_2062 also had high scores in both Δ*ssb* and Δ*dbp*Δ*ssb*. However, proteins with ssDNA binding activities, such as Sul7s (49), RadA (50) and GINS, did not appear in the ssDNA interactome even in the SSB deletion strain (Table S4). The results suggest that these proteins are probably not able to complement the ssDNA-binding function of SisSSB.

**Figure 7.**
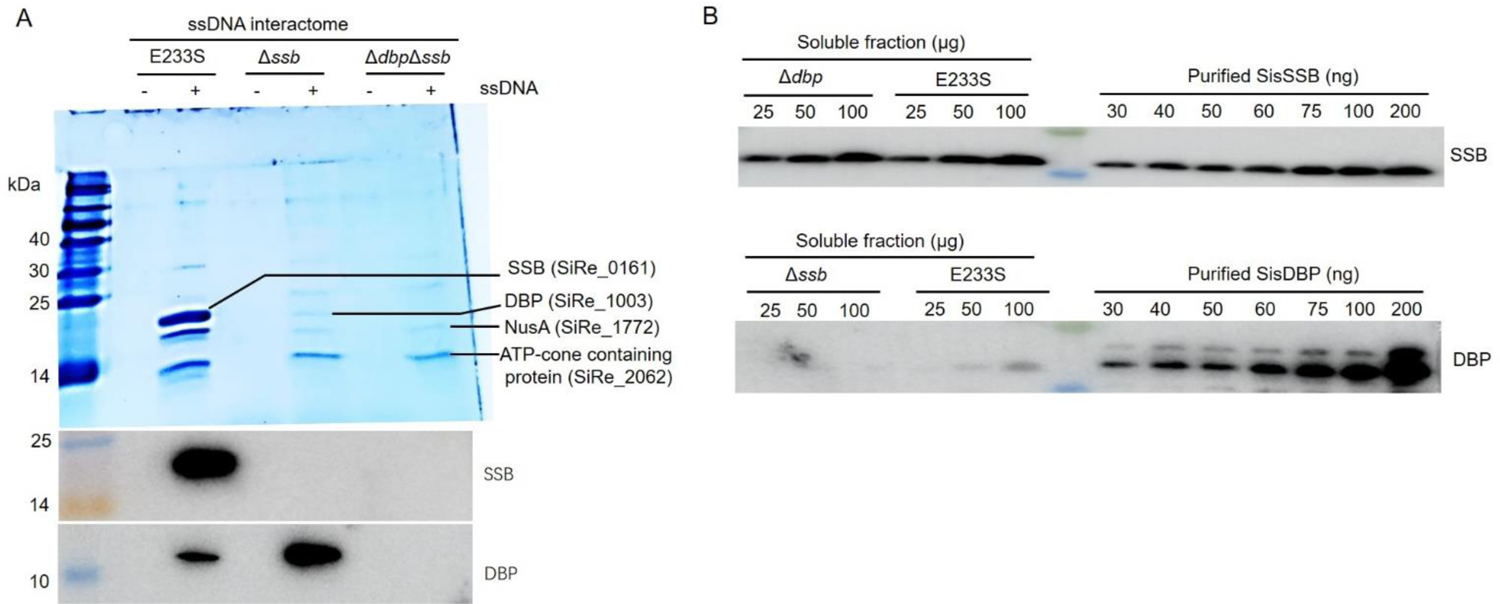
Identification of ssDNA interactome in the cell extract of E233S and estimation of the cellular levels of SisSSB and SisDBP. (**A**) Identification of putative ssDNA-binding proteins using cell extracts of E233S, Δ*ssb* and Δ*dbp*Δ*ssb*. The soluble extracts were separated by SDS-PAGE and analyzed by Western blotting with the antibodies against SisSSB and SisDBP. The gel was stained with Coomassie Brilliant Blue G250. “-”: streptavidin beads without ssDNA; “+”: streptavidin beads with ssDNA. (**B**) Estimation of the cellular levels of SisSSB and SisDBP. The levels of SisSSB and SisDBP in cell lysates were analyzed by SDS/PAGE and Western blotting with SisSSB and SisDBP antibodies. Purified SisSSB and SisDBP in gradient were used for the quantification.

It was estimated that SsoSSB and ThermoDBP accounts for 0.08–0.16% and 0.07– 0.13% of the total soluble protein of *S. solfataricus* and *Thermoproteus tenax* respectively (17). Consistently, SisSSB and SisDBP were estimated as 0.07%∼0.12% and less than 0.01% of the *S. islandicus* proteins, respectively (Figure 7B). The fact that no significant increase of SisDBP in Δ*ssb* suggests that SisDBP is unable to displace SisSSB as an ssDNA binding protein.

### *ssb* deletion did not affect genome stability

To find more evidence that SSB does not participates in DNA replication, genome sequencing of Δ*ssb* and Δ*dbp*Δ*ssb* was performed. The retardant growth of Δ*ssb* at lower temperature could be due to replication defect lacking SSB at the replication fork. In *Saccharomyces cerevisiae*, deletion of Rtt105, a RPA chaperone which facilitates the nuclear localization of RPA and stimulates RPA loading at the replication forks (51), leads to deletions, duplications, and chromosome loss in genome DNA, which resembles replication slippage caused by the absence of pol32 (52). If SSB is involved in replication, growth of Δ*ssb* and Δ*db*pΔ*ssb* at lower temperature may also result in genome instability. Δ*ssb* and Δ*dbp*Δ*ssb* strains were grown at 55℃ for 45 days in liquid STVU medium together with a reference strain Δ*ssb::ssb*. Compare to Δ*ssb::ssb* genome, only few mutations were detected in Δ*ssb* and Δ*dbp*Δ*ssb* genome (Table 2). Both Δ*ssb* and Δ*dbp*Δ*ssb* genomes contain 22 SNPs, but they are not exactly the same (Table 2 and Table S5). For the insertion and deletion of small fragments (<50bp), Δ*ssb* has no insertion and 1 deletion, Δ*dbp*Δ*ssb* has 1 insertion and 2 deletions, one smaller deletion is the same with that in Δ*ssb* (Table S5). No large segment insertion was found in both Δ*ssb* and Δ*dbp*Δ*ssb* genomes, while 1 and 2 large segments deletions were found in Δ*ssb* and Δ*dbp*Δ*ssb* respectively (Table S5) which are engineered gene knockouts. Because there are only few random mutations in Δ*ssb* and Δ*dbp*Δ*ssb* genome (Table 3), we speculate that the occurrence of these variations is independent of the deletion of *Sisssb* and *Sisdbp*. The results indicate that the genome integrity and high-fidelity DNA replication were not compromised in the absence of *ssb* and *dbp*. In addition, flow cytometry of the cells showed that the DNA contents of SSB-deficient strains has no significant difference with E233S or Δ*ssb::ssb* (Figure 2B). These data strongly support that SSB is not a player at the replisome.

**Table 2.** Numbers of mutation events in Δ*ssb* and Δ*dbp*Δ*ssb* at 55℃ by genome sequence *

| | $\Delta ssb$ | $\Delta dbp\Delta ssb$ |
| --- | --- | --- |
| SNP | 22 | 22 |
| Insertion(<50bp) | 0 | 1 |
| Deletion(<50bp) | 1 | 2 |
| Insertion($\geq 50$bp) | 0 | 0 |
| Deletion($\geq 50$bp) | 1 | 2 |

\* *SisSSB in situ* complementary strain $\Delta ssb::ssb$ is used as the reference. SNP: single nucleotide polymorphism.

### The RNA chaperon activity of the SSBs is conserved among archaea which lack Csp homologues

Given that the SisSSB does function in DNA replication but plays a role as an RNA chaperon and Csp, next we want to know whether the RNA chaperon and Csp function is conserved in archaea. For this, we performed bioinformatic, *in vitro* and *in vivo* analyses of SSB proteins from representative archaeal species. Based on amino acid sequence alignment, archaeal single OB-fold SSBs are grouped into three type, SSB-1, SSB-2, and SSB-3 (Figure S6A). SisSSB belongs to the SSB-1 type. We then analyzed the distribution of SSBs/RPA and Csps in the TACK superphyla of Archaea (Figure 8A). Interestingly, most of crenarchaeal species only have one type SSB, SSB-1, and no other SSB/RPA or Csps (Figure 8A). To examine the functions of the SSB1 homologues, we selected four SSB1 homologues (Figure S6A) from species without bacterial Csp homologues (*S. acidocaldarius* DSM 639, *Acidianus hospitalis* W1, *Staphylothermus hellenicus* DSM 12710, *Thermogladius calderae* 1633) and two SSB1 homologues from species having exclusively SSB1 and bacterial Csp homologues (*Candidatus Korarchaeota archaeon* isolate UWMA-0234 and *Candidatus Heimdallarchaeota archaeon* LC_2). As above, pIN plasmids containing these wild type SSBs, SacSSB, AhoSSB, SheSSB, TcaSSB, EcoSSB, KorSSB, HemidallSSB, and their C-terminal truncated mutant SacSSBΔC28, SacSSBΔC51, AhoSSBΔC29, SheSSBΔC17, TcaSSBΔC16, EcoSSBΔC33 (KorSSB and HemidallSSB do not have flexible tail at the C-terminal, Figure S7A) were transformed into *E. coli* BX04 to determine their ability to complement the cold sensitivity of BX04. The results showed that SacSSBΔC51, AhoSSBΔC29, SheSSBΔC17, and TcaSSBΔC16 complemented the cold sensitivity of BX04 as SisSSBΔC29, while KorSSB and HemidallSSB did not, which is similar to that of *E. coli* SSB and *E. coli* SSBΔC33 (Figure 8B and S7B). On the other hand, complementation with full length of SSBs including SacSSB, SheSSB, AhoSSB, and even SacSSBΔC28 is harmful to *E. coli*, similar to that of SisSSB. *In vitro*, SacSSB, AhoSSB, SheSSB, and TcaSSB were able to bind to ssDNA and ssRNA with almost same high affinity, but KorSSB and HemidallSSB have higher binding capacity for ssDNA than ssRNA like EcoSSB (Figure S7B and S7C). Those results suggest that in the archaea containing SSB1 but without bacterial Csp homologue, their SSB1 could act as Csps to destabilize RNA secondary structures at low temperatures and help cells adapt to environment temperature downshift. In contrast, the SSBs possibly lose their RNA chaperone activity when Csp homologous proteins were obtained during evolution.

**Figure 8.**
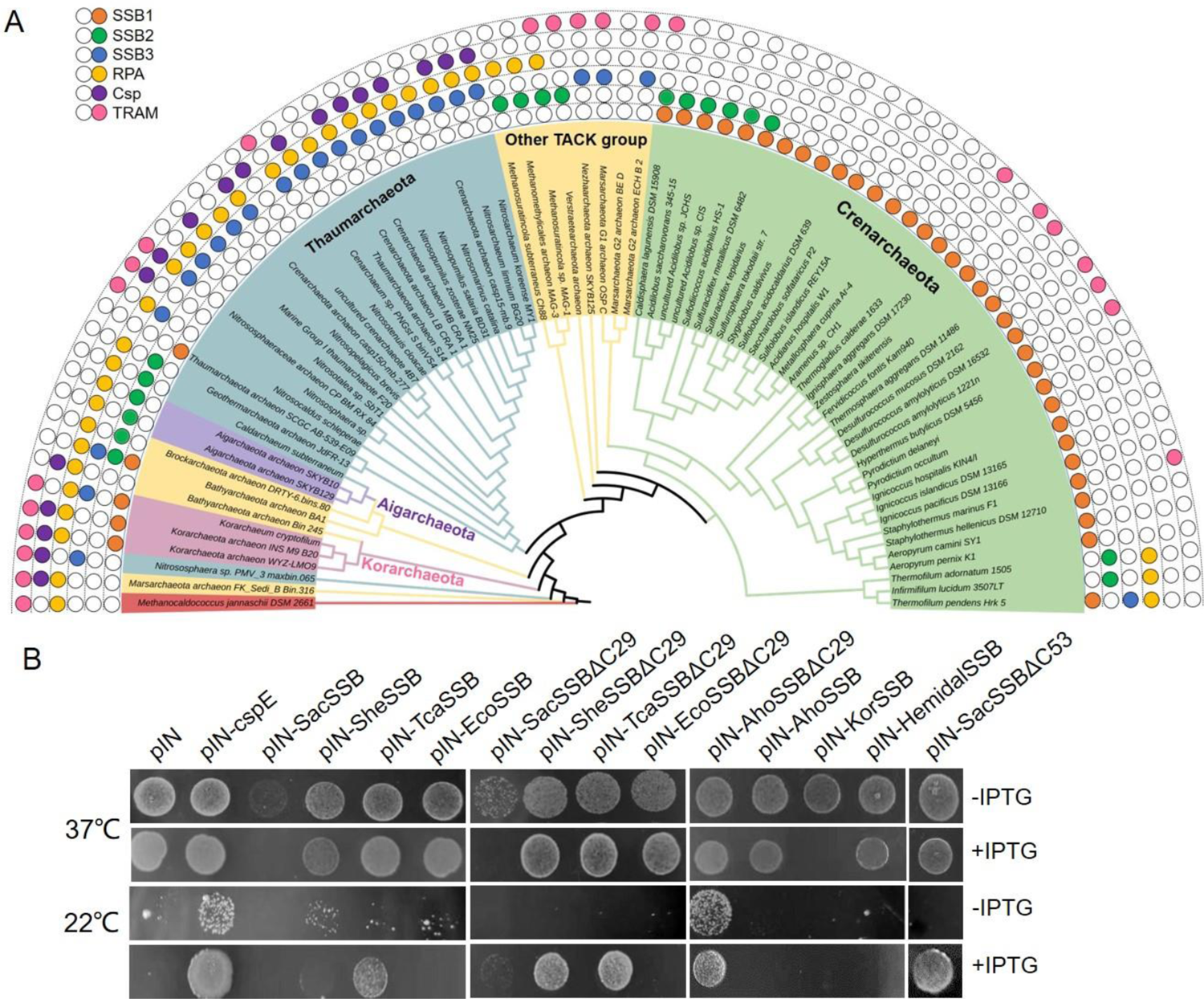
The RNA chaperon activity is conserved among crenarchaeal SSBs. (**A**) Distribution of SSB, RPA, bacterial CSP homologue and TRAM in the TACK superphyla. The phylogenetic tree was constructed using the maximum likelihood method based on 16S rDNA sequences. Totally 77 archaea species were selected in which 76 species were from TACK and *Methanococcus jannaschii* was selected as an outgroup. The bootstrap was 1000. Colored balls indicate presence of the indicated protein while the empty indicates no homology was found. (**B**) The N-terminal OB-fold domains of SheSSB, TcaSSB, SacSSB, and AhoSSB were able to complement the cold sensitivity of *E. coli* BX04.

## Discussion

Although the gene encoding SisSSBs was classified as essential genes in *S. islandicus* species by genome-wide random transposon insertion and identification, it is proven non-essential in this study. The *ssb* knockout strains grow poorly on solid plates by dot assay (Figure S2G) and the colonies of those strains are much smaller than those of E233S (data not shown) when cultured under the same conditions. This may explain the disagreement between our result and result by the genome-wide identification. Our result is in agreement with that by Suzuki and colleague (21) who also found that cells of *S. acidocaldarius* lacking *ssb* was viable and exhibited robust growth in liquid media. Considering the crenarchaeal SSBs displayed similar ssDNA/ssRNA binding activity and *in vivo* phenotypes (Figure 5B and 8B, Figure S5, S6B). We may conclude that all the crenarchaeal SSBs are not essential as a DNA replication and DNA recombination factor. Our transcription analysis of Δ *ssb* indicates that SisSSB may function to protect ssDNA in the cells. Its absence may produce more ssDNA, leading to ignition of the DDR network.

We found one function of SSB is for the cells to adapt to ambient temperature shift. The protein level of SisSSB was up-regulated at lower temperature, which is like the bacterial Csps, albeit with smaller magnitude (Figure 3A and 3B). SisSSB exhibits anti-transcriptional termination ability as Csps (Figure 4A) and is able to complement the loss of Csp in the cold sensitive strain (Figure 5B and Figure S5A). In bacteria, expression of Csps was regulated at the post-transcriptional level through thermosensitive RNA elements, which are always located at the 5′ UTR of mRNA (53,54). While the up-regulation mechanism of SisSSB to adapt temperature downshift needs further exploration.

Sac10b is an abundant protein in *Saccharolobus* cells (55) and its homolog from *S. islandicus* Sis10b was confirmed as RNA chaperone. However, how Sis10b functions *in vivo* is not clear. Genetic analysis suggested that Sis10b is an essential protein, as its knock-down caused reduced growth (40). In our study, SisSSB exhibited a stronger *in vitro* RNA unwinding activity (Figure 4B) and anti-transcription termination in *E. coli* RL211 than Sis10b (Figure 4A). In addition, overexpression of Sis10b did not affect the growth of *E. coli* BX04, and has no help for BX04 to grow at cold temperature. We think that Sis10b functions as a global RNA protector, while one of important role of SisSSB is endowing the cells with adaption to the ambient temperatures downshift.

Archaeal TRAM proteins function as cold shock protein via its RNA chaperone activity (39). However, TRAM is not well-conserved in Crenarchaeota (Figure 8A) and whether TRAM functions as Csp in Crenarchaeota remains unknown. Though TRAM homolog was found in *T. calderae* 1633, TcaSSB binds to ssRNA with high-affinity and the N-terminal of TcaSSB complements the *E. coli* Csps like SisSSB (Figure S6B and Figure 8A and 8B). In addition, bacterial Csp and archaeal TRAM homologs are present simultaneously in some archaea species (Figure 8A), it is not clear how these proteins are functionally distinguished.

An interesting feature for crenarchaeal SSB is that overexpression of SisSSB in *S. islandicus* REY15A caused a significant retardant growth, which is sharply different from bacterial Csps. We found the overexpression of SisSSB affects the cell cycle progression in a global manner (Figure 6D). In addition, SisSSB exhibited higher unwinding capacity of DNA/RNA hybrid (∼47%) than that of dsRNA (∼33%) and dsDNA (∼26%) *in vitro* (Figure S8). A previous investigation showed that SsoSSB had physical and functional interaction with RNA polymerase and it rescues transcription repression by reconstituted chromatin *in vitro* (56). We assume that overexpression of SisSSB leads to a global transcriptional effect which results in retardant growth. However, the exact nucleic acid substrates and the roles of SisSSB in transcription or other processes *in vivo* also needs further investigation.

The levels of bacterial Csps increased from 3-50 times at low temperature (38,54) and decreased after the cells adapted to the low temperature (35,57). In *S. islandicus* REY15A, the SisSSB was up-regulated only about 1.8 times at 55℃. In addition, the overexpression of SisSSB causes serious growth retardance even lethal at normal growth temperature. Thirdly, the basic level of SisSSB in REY15A is high (Figure 7B). These properties of SisSSB differ apparently from those of bacterial Csps which suggests that SisSSB participates other nucleic acid processing in addition to melting RNA secondary structure.

OB-fold containing single-stranded DNA binding proteins play vital roles in DNA replication, HR, DDR, DNA damage repair and genomic stability in almost all living organism (50,58–60). Our genetic analysis reveals that SisSSB plays minor role if any in these processes. A recent study showed that deletion of the gene encoding SacSSB caused strain DP-5 sensitive to DNA damage agents, e.g., cisplatin, metronidazole, and 4-NQO. In addition, the HR frequency of DP-5 was decreased and the loss of SacSSB causes a 29-fold higher mutation rate at low growth temperatures (61). This is different from our study in *S. islandicus* REY15A. After the deletion of *Sisssb*, we could easily knock-out *Sisdbp* coding gene by CRISPR-based genome editing. In this method, the target site was identified and cleaved by the CRISPR-Cas complex in *S. islandicus* REY15A, which causing the genome break and repair by homologous recombination (25,30). The easiness in getting double knockout strain Δ*dbp*Δ*ssb* from Δ*ssb* strain suggests that homologous recombination is independent of SisSSB. The reason behind the differences between ours and the recent report is not clear, it could be due to difference between two microbes. For example, *S. acidocaldarius* differs from *S. islandicus* in several processes, such as cell division (62,63).

Single-stranded DNA binding proteins have been considered as an essential component of DNA replisome (64,65). In addition, the interactome of bacterial SSB and eukaryotic RPA contains numerous DNA replication proteins (8–10,66). Until now, the Crenarchaeota archaeal SSB has only been reported to interact with MCM (mini chromosome maintenance protein) of proteins in the replisome of *S. solfataricus* (67). In our study, we found that SisSSB is not required for the DNA replication in *S. islandicus* REY15A. One possibility is that there is un-identified single-stranded DNA binding protein function in the replisome. Alternatively, the crenarchaeal cell may not require a SSB for DNA replication. Based on the comparison of DNA polymerases, the DNA replication machinery of Crenarchaeota archaea is reported unique (68). Our study may raise an intriguing question about how the crenarchaeal cells maintain faithful replication of the genome in the absence of single-stranded DNA binding protein. Study on the structure and reconstruction of the replisome from crenarchaea would be a promising subject for further investigation.

## Materials and Methods

### Strains and culture conditions

*Saccharolobus islandicus* REY15A(E233S)(Δ*pyrEF*Δ*lacS*) (hereafter E233S) was grown in STVU medium containing mineral salt, 0.2% (w/v) sucrose (S), 0.2% (w/v) tryptone (T), 0.01% (w/v) uracil and a mixed vitamin solution (V). The sucrose (S) is replaced with D-arabinose (A) to make ATVU medium if needed. The medium was adjusted to pH 3.3 with sulfuric acid, as described previously. Normally, cells were cultured aerobically at 75°C with shaking (145 rpm) to OD_600_=0.6-0.8, then transferred to fresh medium with an initial estimated OD_600_ of 0.05 for follow-up growth curve determination.

### Genetic manipulation

The genes encoding SisSSB(SiRe_0161) and SisDBP(SiRe_1003) were knocked out using the endogenous CRISPR-based genome editing system in *S. islandicus* REY15A (25). Firstly, the genomic DNA of E233S was extracted using a bacterial genome rapid extraction kit (SparkJade Co., Shandong, China) and used as the template for high fidelity and SOE (splicing overlap extension) PCR by ApexHF HS DNA polymerase (Accurate Biotechnology Co., Hunan, China) for the donor DNA. The spacer was selected on the target gene consisting of a 5′-CCN-3′ downstream of a 40-nt DNA sequence for making of the artificial CRISPR array in the pGE plasmid. The donor DNA and spacer fragments were ligated to the pGE plasmid with SphI/XhoI and BspQI digestion, respectively. The recombinant pGE plasmid was introduced into the wild type E233S by electroporation. The genotype of single colonies obtained were determined by PCR amplification with Flanking/gene-specific primer pairs. The strains and plasmid constructed and used in this study are listed in Table S1 and the oligonucleotide sequences for PCR and the *in vitro* assay are listed in Table S2.

### Western blotting

Antibodies against SisSSB and SisTBP were produced in rabbit using synthetic specific peptides (amino acids 135-148 RGGRRQENEEGEEE for SSB [SiRe_0161]; amino acids 18–31, SIPNIEYDPDQFPG for TBP [SiRe_1138]). Antibodies against DBP were produced in mouse using the full length DBP expressed in and purified from *E. coli*. To quantify the protein in the cell, the cells (estimated per 5*×*10^8^) were harvested by centrifugation at 5,000 *g* for 10 min and suspended in 100 μL PBS buffer (pH7.4). Then the cells were disrupted by sonication. Total protein was obtained without any centrifugation and the soluble fraction was obtained by collecting the supernatant after centrifugation at 10, 000 *g* for 10 min and filtration with a 0.45 μM cut-off filter. Western blotting was carried out following the standard procedure. The antibodies against SisTBP (loading control) and SisSSB were incubated at the same time.

### Transcriptomic analysis

The transcriptomic analysis was performed as previously described (69,70). E233S, Δ*ssb*, and Δ*dbp* strains were grown at 75℃ with an initial OD_600_ of 0.05 in STVU medium and the cells were harvested when the OD_600_ reached 0.4∼0.5. Cells were pelleted at 6,000 *g* for 10 min and washed in PBS, then frozen with liquid nitrogen and transported with dry ice. The samples were analyzed by Beijing Novogene Bioinformatics Technology Co., Ltd (Beijing, China). Total RNA was extracted using the Trizol reagent (Ambion, Austin, TX, USA) and assessed by Aagilent 2100 bioanalyzer. The libraries were prepared using purified mRNA from total RNA in which the rRNA was removed with probes. The libraries were then sequenced using the Illumina NovaSeq 6000. Filtered high quality clean reads were mapped to the reference genome sequence of *S. islandicus* REY15A (18). Fragments per kilobase of transcript sequence per million base pairs sequenced (FPKM) was used to estimate the gene expression levels, and differential expression analysis (Δ*ssb* versus E233S and Δ*dbp* versus E233S) was performed using the DESeq R package (1.18.0). P values were adjusted using the Benjamini & Hochberg method. Corrected P-value of 0.005 and |log2(foldchange)| > 1 were set as the threshold for significantly differential expression.

### Detection of the ssDNA interactome

The method to detect the ssDNA interactome in *S. islandicus* was according to a previous study (17). Cells were cultured at 75℃ to OD_600_=0.5∼0.8 in STVU medium and collected by centrifugation at 5,000 *g* for 10 min. The pellet was washed with PBS, then resuspended in TEDG buffer (50 mM Tri-HCl pH8.0, 5 mM EDTA Na_2_, 1 mM DTT, 5% glycerol) containing 150 mM NaCl. The cells were disrupted by sonication. After centrifugation (10, 000 *g* for 20 min), the supernatant was filtered with a 0.45 μm cut-off filter. Streptavidin-coated agarose beads 6FF (Smart-Lifesciences, Changzhou, China) were washed three times with NN buffer (20 mM NaH_2_PO_4_, 0.15 M NaCl, pH7.4), then 2.2 nmol (∼31 μg) of 5’ biotined-45nt ssDNA was incubated with the beads with gently shaking at room temperature for 10 min. The beads were washed three times with NN buffer and mixed with 30 mg of the cell lysate. After the mixture was incubated at 55°C for 120 min, the beads were washed 3 times with TEDG buffer containing 150 mM NaCl. Bound proteins were eluted with TEDG buffer containing 1500 mM NaCl. The eluate was TCA/acetone precipitated, resuspended in TEDG buffer, and run on a 15% SDS-PAGE. The gels were stained with Coomassie Brilliant Blue G250 or silver. SisSSB and SisDBP were identified by Western blotting. Distinct protein bands were excised. The excised proteins as well as the whole interactomes were identified by mass spectrometry.

### Flow cytometry

Cells were cultured in liquid STVU medium and estimated 3.0-5.0×10^7^ cells were harvested and fixed with 70% cold ethanol overnight. The fixed cells were pelleted at 4℃, 800 *g* for 20 min. The cells were re-suspended and washed with 1 mL of PBS buffer. The cells were pelleted again, re-suspended in 100 μL of PBS buffer containing 50 μg/mL propidium iodide (PI), and incubated on ice for 30-60 min. The DNA content of the cells was analyzed using the ImageStreamX MarkII Quantitative imaging analysis for flow cytometry system (Merck Millipore, Germany). About 20,000 cells were collected for each sample and the data of the single cells were analyzed with the Flowjo software.

### Cell cycle synchronization

Cells of different strains were synchronized as previously described (26,70). Cells were grown at 75°C with shaking (145 rpm) in 30 mL STVU medium with an initial estimated OD_600_ of 0.045. Acetic acid (6 mM, final concentration) was added when the OD_600_ reached 0.15–0.2 and treated for 6 h. If needed, 0.2% D-arabinose was added after the addition of acetic acid. Then cells were centrifuged at 3,000 *g* for 10 min at room temperature and washed twice with 0.7% (w/v) sucrose to remove the acetic acid. Cells were suspended with 30 mL pre-warmed STVU medium and cultured as above.

### Genome sequencing

Δssb::*ssb*, Δ*ssb* and Δ*ssb*Δ*dbp* strains were first grown at 75℃ and then transferred to 55℃ with an initial OD_600_ of 0.05 for 45 days in liquid STVU medium, during which the culture medium was replaced with fresh medium for three times. The cells were pelleted by centrifugation at 6,000 *g* for 10 min and washed in PBS, frozen with liquid nitrogen and transported under cold conditions with dry ice. Genome sequencing analysis was performed by Novogene (Beijing, China). Briefly, the genomic DNA was extracted and then quantified by Qubit® 2.0 Fluorometer (Thermo Scientific). Sequencing libraries were generated using NEBNext® μLtra™ DNA Library Prep Kit for Illumina (NEB, USA) using 1.0 μg DNA per sample. The whole genome was sequenced using Illumina NovaSeq PE150. The sequenced data were filtered and the clean data were mapped to the reference genome sequence of *S. islandicus* REY15A (18) using BWA software with parameters are as follows: mem -t 4 -k 32 -M -R. SAMTOOLS software was used to count the coverage of the reference sequence to the reads and make explanations of the alignment results following the parameters: depth -d 200000. Single nucleotide polymorphism (SNP) and small fragments (<50bp) variation was e detected by SAMTOOLS. The parameters are as follows: mpileup -m 2 -F 0.002 -d 10000 -u -L10000. Large segments fragments (≥ 50bp) variation found by BreakDancer software. The parameters are as follows: -q 20-d prefix. The obtained variations of Δ*ssb* and Δ*ssb*Δ*dbp* were compared with that of Δ*ssb::ssb* respectively, common mutations are removed and specific variations in Δ*ssb* and Δ*ssb*Δ*dbp* genome were used to show the variations caused at lower temperature.

### Anti-transcription termination assay in *E. coli*

To determine the RNA hairpin melting activity of SisSSB in a heterologous host, a recombinant plasmid pINIII carrying IPTG-inducible SSBs was transformed into *E. coli* RL211. The cells were grown to an OD_600_ of ∼1.0 in LB medium with 100 μg/mL ampicillin. Aliquots (6 μL) of the culture were spotted onto LB plates containing 100 μg/mL ampicillin and 0.2 mM IPTG with or without 30 μg/mL chloramphenicol. The plates were incubated at 37℃ for 2–3 days. *E. coli* RL211 carrying IPTG-inducible CspA, CspE or Sis10b or with an empty pINIII were used as positive and negative controls, respectively.

### Protein purification

Gene fragments of SisSSB, SacSSB, SheSSB, TcaSSB, and EcoSSB and their mutants were obtained by PCR. Gene fragments of AhoSSB, KorSSB, HemidallSSB, and KorSSB were obtained by chemical synthesis after codon optimization. The SSB genes were then cloned into pET15bm plasmids with *Nde*I and *Sal*I restriction sites and transformed into *E. coli* BL21(DE3) CondonPlus-RIL. Cells were cultured at 37℃ with 180 rpm shaking to OD_600_=0.4∼0.6 in LB medium with 100 μg/mL ampicillin sodium and 34 μg/mL chloramphenicol. IPTG (0.3 mM) was added and the cells were cultured at 37℃ for 5 h. The cells were harvested by centrifugation and resuspended in BufferA (50 mM Tris-HCl, pH7.4, 200 mM NaCl, 5% glycerol) and lysed by sonication. The cells carrying no tag SisSSB, SacSSB, SheSSB, TcaSSB, and AhoSSB were disrupted by sonication. Cell extracts were heated for 20 min at 65°C and then centrifugation at 4℃, 10, 000 rpm for 20 min. The cell debris of strains carrying plasmid expressing C-His-tagged KorSSB, HemidallSSB, and Eco SSB was removed by centrifugation (without heat treatment). Polyethyleneimine (PEI) (0.5%, W/V) was added into the supernatant to remove the nucleic acids. The mixture was centrifuged at 10,000 *g* for 30 min at 4℃. The proteins were then precipitated with 0.6 g/mL (NH_4_)_2_SO_4_ and collected by centrifugation under the conditions as above.

The protein samples were then dissolved and dialyzed overnight in buffer A. The dialyzed mixture was centrifuged, and the supernatant was then filtered through a membrane filter (0.45 μm). The non-tagged proteins were purified by a cation exchange column (SP) and a Heparin column sequentially. The proteins were eluted in buffer B (50 mM Tris-HCl pH7.4, 1.5 M NaCl, 5% glycerol) with 0-100% salt gradience. The fractions were collected at 50%-70% salt gradience. The C-his SSBs samples were load into a Ni column pre-equilibrated with buffer A and the proteins were eluted in buffer A with 400 mM imidazole. For further purification, proteins were loaded into a size exclusion chromatography column Superdex 200 increase 10/300 (GE Healthcare). The protein concentration was determined by the Bradford method.

### RNA unwinding assay

A 40 nt ssRNA FAM-labelled at the 5′ end was annealed to a 40 nt partial complementary ssRNA labelled at the 3′ end with a BHQ1 quenching at a molecular ratio of 1:1. Annealed dsRNA products were 22 bp paired fragments (*T*m: 59°C) at one end and 18 bp unpaired bases at another end. The wild-type SisSSB (20 μM) was incubated for 30 min at 37℃ with 100 nM annealed dsRNA probe in 200 mM Tris– HCl, pH 7.4, and 10 mM MgCl_2_. Changes in fluorescence values were measured using a fluorescence spectrophotometer (ENSPIE-2300, PE, America) with a 460 nm excitation wavelength and a 515 nm emission wavelength.

### Complementation of the cold-sensitive strain *E. coli* BX04

SSBs and their mutants were cloned into pINIII plasmids with *Nde*I and *Bam*HI restriction sites and transformed into *E. coli* BX04. Colonies were picked and cultured overnight in LB medium with 100 μg/mL ampicillin under 180 rpm shaking at 37℃. The overnight culture was transferred to fresh LB medium containing 100 μg/mL ampicillin with 1% inoculum and incubated for 3−5 h. Cultures were adjusted to OD_600_ 1.0 with fresh medium, diluted and spot on the LB plates with 100 μg/mL ampicillin, with or without 0.2 mM IPTG then incubated for 2-5 days at 37℃ and 22℃, respectively.

### Electrophoretic mobility shift assay (EMSA)

The 40 nt 5’-FAM-labelled ssDNA (ssDNA-FAM) and ssRNA (ssDNA-FAM) were used as the substrates to determine the binding capacity of SSBs. The binding assay was performed in a 20 μL reaction mixture containing 20 nM ssDNA/ssRNA, 50 mM Tris-HCl, pH 7.4, 5 mM MgCl_2_, 20 mM NaCl, 50 μg/mL BSA, 1 mM DTT, 5% glycerol, and different concentrations of purified proteins (for ssRNA, 0.5 U/μL RNase inhibitor was added). The reaction mixture was incubated at 37°C for 30 min before loaded onto a 12% native polyacrylamide gel. After running in 0.5×TBE, the gel was visualized using an Amersham ImageQuant 800 biomolecular imager (Cytiva).

## Acknowledgements

This work was supported by the National Key Research and Development Program of China (No. 2020YFA0906800), the National Natural Science Foundation of China (No. 31970546 and 31670061 to YS, 31970119 to JN, and 31771380 to QS) and the State Key Laboratory of Microbial Technology. We would like to thank Prof. Li Huang for providing bacterial strain and plasmid of Sis10b for the analysis. We also thank all the lab members for helpful discussions. We thank Dr. Jiangchuan Shen at Indiana University for stimulating discussions.

## Author contributions

Y.X., Q.H. and Y.S. conceived and designed the research, and prepared this manuscript; Y.X. and Z.J. carried out experimental work; M.Z. and P.W. helped bioinformatic analysis; X.Z. helped proteins purification and EMSA assay; Q.G. and Y.Y. helped cell cycle synchronization and data analysis; Q.S. advised ssDNA interactome detection; X.F. advised genome analysis and helped RNA assay; J.N. helped the transcriptomic data analysis; X.D. helped cold sensitive assay. All authors edited the manuscript.

## DATA AVAILABILITY

All data supporting the findings of this study are available within the article and its Supplementary Information, or from the corresponding author upon reasonable request. The genome DNA sequences and RNA-seq data were deposited under the accession number PRJNA982601 and GSE234768, respectively.

